# Residence Times and Legacy of Biogenic Carbon in Ocean Reservoirs

**DOI:** 10.1101/2024.09.06.611583

**Authors:** André W. Visser

## Abstract

Quantifying the sequestration potential of biologically driven carbon fluxes in the ocean depends critically on residence times – how long carbon remains stored in reservoirs before being re-exposed to the atmosphere. Simple mass balance provides estimates for many of the major ocean biogenic carbon reservoirs. For vegetated coastal ecosystems (mangroves, sea grass meadows, salt marshes) that globally store 20 to 40 PgC, this is 200 to 500 years, while for the biological carbon pump, a reservoir of about 2000 PgC, it is between 200 to 800 years. Over these time scales respective reservoirs reach equilibrium if left undisturbed. Importantly, near equilibrium of ocean reservoirs during the Holocene can be inferred from the near steady atmospheric concentrations during this period. The degradation of habitats and the over-exploitation of living marine resources particularly in the last 75 years have tipped these natural processes out of balance, to the extent where many are now net emitters of legacy carbon back to the atmosphere. The analysis exposes a conflict between how sequestration is reported in oceanographic literature and how it is understood with regards durable carbon capture and storage. Nature-based solutions can be sought to address parts of the climate crisis, by improving ecosystem health and biodiversity, but are unlikely to provide solutions to carbon management on a scale commensurate with anthropogenic emissions. The best we can do is to limit net emissions by restoring what we can, and to ensure that future practices do not further tip ocean carbon reservoirs out of balance.

**Significance Statement:** Marine animals and plants maintain large pools of carbon in the ocean and coastal areas that have been laid down by generations past. This legacy carbon is continuously being recycled on time scales of 100s of years. Left undisturbed, as they were for most of the last 10000 years, these carbon pools tend to equilibrium; flux in equals flux out. Human activities such as over fishing and coastal construction, particularly in the past 75 years, have tipped these natural cycles out of balance to the extent where many pools are now net emitters of carbon. Conservation and restoration of marine habitats can bring these cycles back into balance but cannot be counted as offsetting fossil fuel emissions.

---

A significant fraction of the carbon in the Earth System is stored in the oceans. This is second only to the lithosphere, and considerably more (by a factor of over 40) than the carbon currently in the atmosphere. This observation has focused considerable attention on ocean carbon cycles and budgets, largely with a view to documenting the role of the oceans in regulating global climate and its potential in soaking up anthropogenic carbon emissions (1–4). While it ‘s true that the oceans have taken up about 30% of emissions through physics and chemistry in equilibrating with increased atmospheric partial pressure, a cursory examination of current oceanographic literature would leave the impression that oceans and coastal regions are also remarkably adept at sequestering carbon via biological processes, for instance through the biological carbon pump (5–8) or in coastal vegetated ecosystems (9–12). This has prompted suggestions that nature-based solutions to our climate crisis are not only effective (13, 14), but valuable trading commodities (15), a theme that is echoed in numerous governmental and NGO websites, media outlets and commercial prospectuses. But how well are such conclusions founded? In particular, are we as marine scientists and oceanographers being less than rigorous when it comes to communicating and self-critiquing what we actually mean by carbon sequestration, and does this correspond to how it is generally understood in the broader context of carbon capture and storage (CCS)?

Carbon stored in the Earth System can be broken down into different reservoirs (16) depending on province and provenance. Figure 1 outlines our current understanding of the size of these reservoirs with respect to the oceans. The largest is the dissolved organic carbon (DIC) reservoir which is estimated at 38,000 PgC. While most of this DIC reservoir is attributed to physicochemical processes – Henry ‘s law and carbonate chemistry – a significant fraction is maintained by the biological carbon pump (BCP) (17). Recent estimates (3, 18) place BCP contributions to the DIC ocean reservoir at about 2000 PgC via the respiration of organic material (the soft tissue pump) and about 800 PgC via biogenic carbonates. Other significant oceanic carbon reservoirs include about 700 PgC dissolved organic carbon (DOC) (19) and the 2200 PgC in various forms of carbon found in marine sediments (20). A final reservoir to note is that associated with vegetated coastal ecosystems – often referred to as blue carbon – totalling between 10 to 24 PgC (10, 21). The respired DIC reservoir, the DOC reservoir, the biogenic carbonate reservoir and some part of the marine sediment reservoir are biogenic in origin, and can be attributed to the various pathways of the biological carbon pump (5). It is of some interest to note that the sum of these biogenic carbon reservoirs, is about 3500 PgC and far exceeds the total living biomass of the oceans, estimated at about 3 to 5 PgC (22). This includes everything from microbes to whales. Seen in this light, the carbon capital of marine organisms is not so much in their living biomass, but rather in the legacy carbon laid down by preceding generations.

**Fig. 1.**
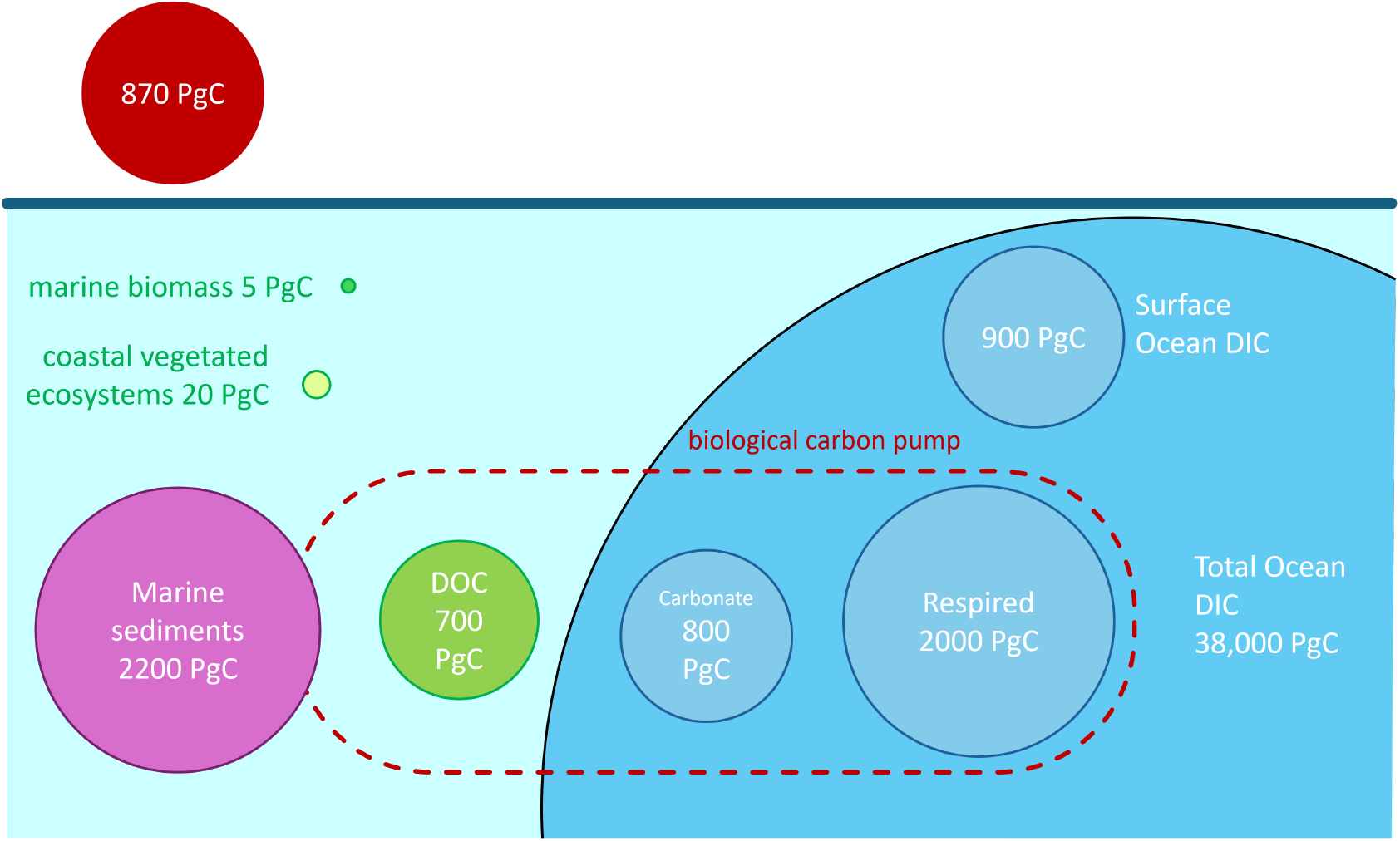
Estimated size of oceanic carbon reservoirs. The size of the bubbles (in area) is proportional to the size of the reservoirs.

This system of interconnected reservoirs is dynamic in two distinct ways. Firstly, carbon is continuously being cycled between reservoirs with characteristic residence times in each, and secondly the various reservoirs themselves can expand or contract. In terms of carbon and climate, the relevant dynamic is associated with the expansion or contraction of reservoirs. This is precisely what is inferred when we speak of carbon sequestration. Namely, are any of the ocean reservoirs expanding to offset some of the 10 PgC anthropogenic emissions released to the atmosphere annually? Perhaps even more important, are there other anthropogenic practices (e.g. fishing, dredging, coastal construction) that are causing reservoirs to shrink releasing legacy carbon to the atmosphere?

We can make this a bit more concrete by considering the mass balance of a reservoir. Specifically, at steady state, the residence time *T* [yr] of carbon in a reservoir, the turn-over flux *Q* [PgC yr^−1^] and the mass of a reservoir *B* [PgC] are related as

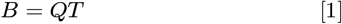

(cf Methods). The dynamics here represent the more rapid turn-over of the reservoir. We can define sequestration in the sense of a potential offset to anthropogenic emissions as any process that leads to an increase in *B*, i.e. *dB/dt >* 0. There is of course also the possibility that *dB/dt* < 0 indicating a net efflux of carbon out of the prescribed reservoir. Seen in this sense, sequestration arises though either an increase of flux into a reservoir, or an increase of residence time. Notably, sequestration is not equivalent to *Q*, even if it represents carbon that will not return to the atmosphere for a geologically significant time scale.

To underscore this, Figure 2 shows the atmospheric CO_2_ concentrations over the Holocene (23, 24), a period of extremely stable climate following the last glacial maximum about 12,000 years ago. During this time, atmospheric CO_2_ remained stable at about 280 ppmv, and there is no reason to believe that there were any major changes in how the natural carbon cycles of the Earth System operated then and now. Plankton communities flourished and rained detrital material into the ocean ‘s depths, large marine metazoans migrated and respired and their dead bodies sank to the sea floor, vegetated coastal ecosystems accumulated organic rich material into their sediments, and the meridonal overturning circulation steadily stirred the oceans from top to bottom. Indeed, if we were to look to a time in the earth ‘s geologic past that best typified the pristine baseline natural state to which conservation and restoration efforts aspire, then the mid Holocene would be very close to the mark. This relatively unperturbed state of the Earth System remained in effect until the great acceleration of the Anthropocene from about 1950 on (25).

**Fig. 2.**
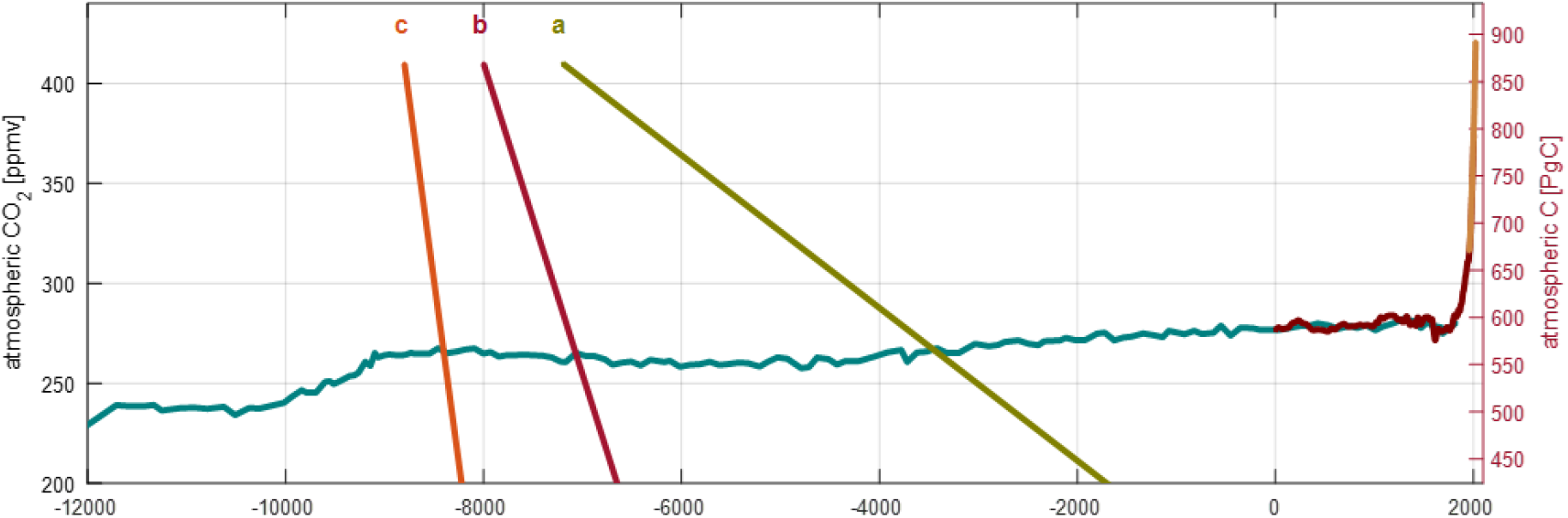
Atmospheric carbon reservoir during the Holocene. Conversion from ppmv (parts per million by volume) CO2 to carbon content follows the IPCC with a conversion factor of 2.12 PgC/ppmv. The corresponding atmospheric reservoir mass in PgC is qiven on the right axis. Superimposed are sequestration rates suggested in the literature: (a) 81MtC /year for vegetated coastal ecosystems (15), (b) 330 MtC yr^−1^ and (c) 720 MtC yr^−1^, the lower and upper estimates for the biological carbon pump suggested by (26). The higher estimates suggested by (7) are even steeper and is not included here.

In Figure 2, in addition to the observed atmospheric concentrations, I have also plotted the sequestration rates reported in various contexts in the literature. For instance, vegetated coastal ecosystems are reported to sequester in the order of 80 MtC yr^−1^ (15) while estimates for the biological carbon pump range from 720 MtC yr^−1^ (8) to considerably higher 900 to 2600 MtC yr^−1^ (7). No sequestration, in the sense of a durable drawing down atmospheric CO_2_ at anywhere near these rates is apparent. Even the relatively gentle sequestration rate for vegetated coastal ecosystem much heralded in blue carbon literature (13, 15) would have emptied the atmosphere of CO_2_ in about 5000 years. Clearly this didn ‘t happen. In the absence of any evidence indicating global carbon cycles were radically different, this would suggest that during the bulk of the Holocene, oceanic reservoirs were in near equilibrium (27). The rates suggested here cannot be equated with sequestration in the sense of CCS, rather they are more likely to represent the turn-over fluxes *Q* associated with their respective reservoirs.

Indeed, if we consider these as turnover-fluxes, then by Eq(1) we can estimate, at least in an order of magnitude sense, the respective residence times for various reservoirs. For instance, for vegetated coastal ecosystems, *B* is in the range 10 to 24 PgC, and *Q* approximately 80MtC yr^−1^ suggests a residence time would be a few hundred years. For the 2000 PgC associated with respired organic carbon with *Q* in the range 5 to 10 PgC yr^−1^, the residence time would be of a similar scale (cf Methods). The picture that emerges is that major biogenic oceanic carbon reservoirs became relatively stable within a 1000 years or so of the end of the last glaciation, after which they were maintained in equilibrium by the accumulation of new carbon to replace the release of legacy carbon laid down by preceding generations of biota.

“Every year, coastal wetlands sequester enough CO_2_ to offset the burning of 1 billion barrels of oil “ reads a promotional flyer published by a well-intentioned conservancy group. Skimming through scientific literature, it is easy to see where this and similar statements arise. At the heart of this mis-communication is that the marine science and oceanographic community has lifted the definition of sequestration directly from carbon capture and storage (CCS), namely that carbon can be deemed to be sequestered if it is removed from the atmosphere for more than 100 years (28–32). This definition works well enough when speaking of artificially capturing carbon and sticking it in a hole in the ground. For natural systems such as vegetated coastal ecosystems and the biological carbon pump that have been in existence longer than their residence times, this definition has little meaning: most of the carbon entering these reservoirs is simply replacing that which is leaking out. Technically speaking, the CCS definition of sequestration, whether it is based on 100 year or indeed any time horizon, is a necessary but not sufficient condition for what can be understood as durable carbon sequestration.

The concept of a minimum time sequestration time scale is intuitive but ultimately not particularly helpful. It is somewhat ironic for instance, that the one oceanic reservoir that has perhaps been most effective in the uptake of anthropogenic C0_2_ emissions, namely the surface ocean, is often discounted as a site for carbon sequestration because of its high turnover rate with the atmosphere. This is rapid, on the scale of months to years depending of depth and location, but its sheer volume and absorption capacity means that the surface ocean has made a serious dent (33–35) in the approximately 450 GtC that has been released to the atmosphere from fossil fuels since the industrial revolution (27). Large turnover rates even with short residence times can still maintain reservoirs of significant size.

Understanding the oceans ‘ role in climate really boils down to describing and predicting the expansion or contraction of the major carbon reservoirs of marine systems. Of particular interest are any systematic changes in processes that impact both the turnover flux and the residence time. As a case in point, predicting the expected response of the biological pump to climate change has focused largely on export flux *Q*, where best estimates indicate a ∼10% reduction on a global scale (36). However, the process that leads to this reduction - an increase in ocean stratification - also results in a more sluggish meridional overturning circulation (37) meaning that what goes down, stays down longer i.e. longer residence times. Simulations indicate that increased residence times wins out over decreased export flux so that the net effect is an increase in the size of the respired DIC reservoir in the range 20 to 50 PgC by the end of the 21st century (38, 39), i.e. the biological carbon pump may store more carbon under climate change providing a negative feedback. It can be conjectured that ultimately, it is the more rapid remineralization and injection of nutrient compared to respired DIC that drives this trend.

## Conclusions

The worlds oceans and marine environments maintain climatically significant stocks of biogenic carbon that have been laid down by generations of biota spanning back 100s to 1000s of years. It appears that during the past 10,000 years, these stocks were in near equilibrium with fluxes in balancing fluxes out, a situation that persisted up until about 75 years ago. Whatever changes have occurred in the Earth System since then, there are still vast pools of legacy biogenic carbon in the oceans and marine environments (∼3500 PgC) that are inexorably being recycled and re-emitted to the atmosphere. Without healthy and fully functioning ecosystems to maintain the equilibrium, the oceans and coastal seas have become, and will continue to be net emitters of biogenic carbon to the atmosphere; a point already noted for various biogenic carbon reservoirs. Examples include the emissions incurred by the rapid loss of vegetated coastal ecosystems over recent decades (40), the intensified harvesting of targeted fisheries over the past 40 years (46), and the lingering effects of the decimation of baleen whale populations in the 1960s (47). Conservation and restoration efforts are vital for maintaining legacy carbon reservoirs as well as improving ecosystem health and biodiversity and providing coastal protection against sealevel rise, but cannot be used to offset fossil fuel emissions; any attempt to do so should be called out as false accounting. Novel marine ecosystems, whether artificially constructed, or emerging through, for instance, disappearing sea-ice in high latitude seas and coastal areas, could provide some additional sequestration. But even if there were a doubling of vegetated coastal ecosystems world-wide, this would only offset a few years of fossil fuel emissions at current rates. The use of the oceans and marine environments to manage global carbon will always be fraught with two insurmountable problems: (1) under natural conditions, even returning to the pristine state of the mid-Holocene, the potential for expanding biogenic carbon reservoirs is simply too small to make a difference, and (2) if scaled up to make a difference, the solution can no-longer be deemed natural with serious consequences for the ocean ‘s bio-geochemical cycles and marine life (48).

In addressing ocean carbon budgets and climate, it is vitally important to be mindful of the legacy of biological processes in bygone ages. Failure to do so leaves our estimates of global carbon budgets unfit for purpose. Hyperbolic language is not only unhelpful, it is detrimental to an informed political solution to our climate crisis. Sequestration is a term that carries a lot of weight in this discourse, and should be avoided unless it refers to a net mass balance gain for a non-atmospheric reservoir. We should also be mindful of our footprint on natural cycles involving biogenic carbon. The harvesting of marine living resources and the conversion of coastal habitats incur costs in legacy carbon emissions that are poorly understood, but should be accounted for.

## Materials and Methods

### Vegetated Coastal Ecosystems

Numerous studies have provided estimates of carbon stocks and accumulation rates for vegetated coastal ecosystems (9, 10, 41). These estimates largely agree on the carbon stock density in the range 140 to 280 MgC ha^−1^ (specifically, accumulated organic carbon the upper 1 m of sediment), with mangroves tending to the higher end, and sea grass meadows to the lower (Table 1). Averaged accumulation rates are also relatively consistent ranging from 80 to 240 MgC km^−2^ yr^−1^. When integrating these estimates up to a global scale, the largest uncertainty appears in the areal extent of these habitats, which yield global stock estimates in the range 7 to 23 Pg C, and accumulation rates ranging from 40 to 200 TgC yr^−1^. These stock estimates however, only represent the upper 1 m of sediments that can extend to several meters thickness (**?**). Taking a mean depth of 3 m and a global surface storage density of about 13 PgC gives a total carbon stock of about 40 Pg for vegetated coastal ecosystems. Likewise, a relatively conservative global averaged accumulation rate would be about 88 TgC yr^−1^ (15), a figure that is also lent credence given the statistical bias of current estimates (44, 49). This yields a residence time estimate of *T* = *B/Q* ∼400 years (range 200 - 500 years). A similar number is arrived at bypassing the uncertainty in areal extent. While there remains a large uncertainty in this estimate, largely through our assumed accumulation thickness, it is consistent with observed remineralization rates. Specifically, under anoxic conditions as is found in the sedimentary deposits of many vegetated coastal ecosystems the dissolution of some of the more refractory components of solid organic material such as lignins have half lives of a few 100 years (50).

**Table 1.** Stocks, accumulation rates and residence time estimates for vegetated coastal ecosystems.

| Habitat | $\rho$<br>[MgC ha<sup>-1</sup>]<br>(40) (10, 41) | | $\Omega$<br>[1000 km<sup>2</sup>]<br>(10) | $C$<br>[PgC]<br>$C \approx \rho\Omega$ | $\alpha$<br>[MgC km<sup>-2</sup> yr<sup>-1</sup>]<br>(41–45) | | $A$<br>[TgC yr<sup>-1</sup>]<br>$A \approx \alpha\Omega$ | $T$<br>[yr]<br>$T \approx hC/A$ $T \approx h\rho/a$ | |
| --- | --- | --- | --- | --- | --- | --- | --- | --- | --- |
| Salt marshes | 250 | 160 | 200<sup>400</sup><sub>20</sub> | 4<sup>10</sup><sub>0.3</sub> | 240 | 220 | 46<sup>96</sup><sub>4</sub> | 270<sup>310</sup><sub>220</sub> | 270<sup>340</sup><sub>200</sub> |
| Mangroves | 280 | 260 | 140<sup>150</sup><sub>130</sub> | 3.8<sup>4</sup><sub>3</sub> | 170 | 160 | 23<sup>26</sup><sub>21</sub> | 490<sup>490</sup><sub>480</sub> | 490<sup>530</sup><sub>460</sub> |
| Sea grass | 140 | 140 | 400<sup>600</sup><sub>180</sub> | 5.6<sup>8</sup><sub>2</sub> | 80 | 140 | 44<sup>84</sup><sub>14</sub> | 380<sup>530</sup><sub>300</sub> | 410<sup>530</sup><sub>300</sub> |
| Total |  |  |  | 13<sup>23</sup><sub>6</sub> |  |  | 110<sup>210</sup><sub>40</sub> | 360<sup>530</sup><sub>220</sub> | 390<sup>450</sup><sub>330</sub> |

Carbon density $\rho$ [MgC ha<sup>-1</sup>] and global carbon stocks $C$ [PgC] only include the upper 1 m of sediments. To get the integrated stock, I assume a sediment thickness of $h = 3$ m. The areal extent of habitats $\Omega$ [1000 km<sup>2</sup>] exhibit a range, where the subscript and superscript on the mean estimate refer to lower and upper bounds(10). Mean accumulation rate $\alpha$ [MgC km<sup>2</sup> yr<sup>-1</sup>] come from various referenced sources, but appear relatively well constrained. Global accumulation rates $A$ [TgC yr<sup>-1</sup>] exhibit a range largely due to the variation in areal extent estimates.

### The Biological Carbon Pump

The biological pump is evidenced by the vertical gradient of DIC observed in the ocean. In part, this gradient is maintained by the respiration of organic matter originating from the surface ocean and transported to depth either as sinking detrital material or vertically migrating organisms. How much carbon is stored in the oceans through the oceanic carbon pumps can be deduced from the distribution of chemical tracers. Integrating the observed distributions of DIC yields a global inventory of 38,000 PgC in the oceans. A simple examination of observed gradients suggests that about 10% (i.e. ∼4000 PgC) is involved in the various oceanic carbon pumps. Of this about 2/3 (3, 51) is due to the biological pump (i.e. ∼2600 PgC). It is also evident from DIC profiles that the bulk of the stored carbon (*>* 80%) is below 1000 m depth (i.e. ∼2100 PgC).

More recently, an inverse biogeochemical model (18) based on ocean circulation, a bulk uniform Redfield ratio and observed distributions of tracers (DIC, alkalinity, nutrients, oxygen) provided estimates of 1300 PgC due to the respiration of organic matter, and 540 PgC due to the dissolution of biogenic carbonates. Similar estimates were arrived at using a global food-web model (52). As pointed out by (3), these are underestimates as they assumed a near instantaneous equilibrium between the atmosphere and surface ocean, a process that is actually quite sluggish with time scales of about 1 year. Specifically, these estimates assume the residence time to be the same as the mean first passage time (53) – i.e. the average time between the injection of DIC at some depth and location and when that DIC first encounters the ocean surface. A significant fraction of this DIC is subducted as preformed DIC before complete equilibrium can be effected, meaning that the actual residence time is significantly longer that the first passage time. Correcting for this, (3) estimated an additional 1000 PgC of biogenic DIC in the global inventory, which translated roughly into totals of 2000 PgC due to the respiration of organic matter, and 800 PgC due to the dissolution of biogenic carbonates.

The biological carbon pump is ultimately controlled by export production, the amount of biogenic carbon that is exported from the surface ocean (i.e. below the euphotic zone 100m) to the ocean ‘s interior. While there is a relatively broad range of estimates, many cluster around 10 PgC yr-1 (54, 55), and a fairly simple estimate of the mean residence time for respired carbon is thus 200 years (52). Estimates of flux through the permanent thermocline 2 PgC/year maintaining about 80% if the respired DIC reservoir would suggest a residence time of about 800 years. A rough estimate for the residence time for respired DIC is thus ∼500 years (range 200 to 800 years), which is also consistent with more precise residence times estimated for specific pathways of the biological pump (Table 2).

**Table 2.** Estimates of reservoir mass *B*^∗^ [PgC] and first passage time estimates *T* ^∗^ [years] for pathways in the BCP.

| | $B^*$ | $T^*$ |
| --- | --- | --- |
| Export Flux (52) |  |  |
| –Phyto detritus | 210<sup>300</sup><sub>100</sub> | 180<sup>200</sup><sub>170</sub> |
| –Fecal pellet | 830<sup>950</sup><sub>750</sub> | 130<sup>140</sup><sub>130</sub> |
| –DVM | 150<sup>190</sup><sub>80</sub> | 150<sup>200</sup><sub>100</sub> |
| –Mixing | 102<sup>105</sup><sub>100</sub> | 50<sup>51</sup><sub>49</sub> |
| Total | 1290<sup>1540</sup><sub>1030</sub> | 130<sup>135</sup><sub>125</sub> |
| Metazoans Vinogradov's ladder (56) |  |  |
| –Mesozooplankton | 120<sup>270</sup><sub>40</sub> | 180<sup>410</sup><sub>70</sub> |
| –Macrozooplankton | 230<sup>380</sup><sub>110</sub> | 210<sup>350</sup><sub>110</sub> |
| –Mesopelagic | 280<sup>710</sup><sub>50</sub> | 270<sup>500</sup><sub>160</sub> |
| –Forage fish | 70<sup>110</sup><sub>30</sub> | 470<sup>560</sup><sub>260</sub> |
| –Large pelagic | 20<sup>40</sup><sub>5</sub> | 410<sup>480</sup><sub>340</sub> |
| –Gelatinous zooplankton | 70<sup>120</sup><sub>30</sub> | 640<sup>870</sup><sub>490</sub> |
| Total | 780<sup>1200</sup><sub>480</sub> | 250<sup>390</sup><sub>150</sub> |
| From estimated abundance and distribution |  |  |
| Overwintering copepods (57) | 14 – 37 | 160 – 750 |
| Krill fecal pellets southern ocean (31) | 9 – 2 | 200 – 60 |
| Cephalopod deadfall (58) | 0.002 – 0.005 | 200 – 1000 |

These estimates are based on first passage times. Accounting for preformed fractions, total reservoir mass can be estimated as $B = 1.6 \times B^*$ and mean residence time as $T = 1.6 \times T^*$ (Methods)

### Reservoir Mass Balance and Characteristics

Carbon reservoirs can be defined in terms of their province and provenance (where it is and how it got there). Mathematically speaking they are subsets of all the carbon in the Earth System. Definitions need not be unique, a reservoir can be a subset of another and reservoirs can intersect. It is no accident that Fig 1 resembles a Venn diagram unsed to illustrate logical relationships in set theory. Seen in this light, sequestered carbon simply all carbon in the Earth System that is not in the atmosphere. More importantly, the definition of reservoirs is such that it ensures mass conservation, and the mass of a reservoir can only change by changes in what comes in or what goes out.

Any given reservoir, irrespective of the details of its dynamics, can be defined in terms of residence time weighted influx and efflux functions, *q*(*θ, t*) and *p*(*θ, t*) . Specifically *q*(*θ, t*)*dθ* is the influx of material at time *t* that will remain in the reservoir for a period in the range [*θ, θ* + *dθ*]. Similarly, *p*(*θ, t*)*dθ* is the efflux of material at time *t* that has had a residence time in the reservoir in the range [*θ, θ* + *dθ*]. These functions, although in general different, are not independent. Conservation of mass within the reservoir means that the efflux of material at time *t* that had residence time *θ* must be equal to the influx with the same residence time that entered the reservoir at time *t* − *θ*. That is

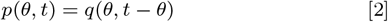

By symmetry, this also means that

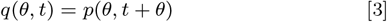

These are general conditions for any mass conserving system, and a more rigorous mathematical approach can be taken in terms of the McKendrick-von Foerster equation (59). Further, these conditions are not dependent on the internal dynamic of the reservoir, whether it is composed of multiple interconnected compartments, and/or the dynamics are time dependent (60, 61). Such facets will change the form of *q* and *p* but not their dependence expressed in Eqs (2) and (3). The total influx at time *t* is

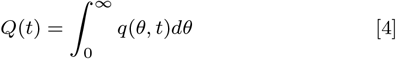

and likewise

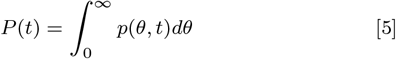

is the total efflux at time *t*. Noting that *p*(*θ, t*) contains all the information on residence times prior to *t*, while *q*(*θ, t*) describes residence times after *t*, the total mass of the reservoir can also be derived as

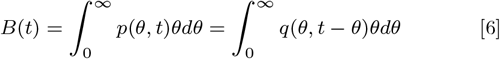

Finally, the mean residence time can be formulated as

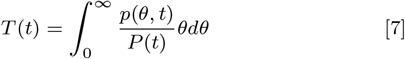

Or more succinctly as

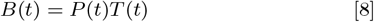

This relationship is quite general and describe in a rigorous sense how the mass of a reservoir changes in terms of total efflux *P* (*t*) and mean residence time *T* (*t*). In a practical sense however this is not super useful as it stands. Estimating *q* and *p* is far from trival, and very few studies have focused on the efflux from earth system carbon reservoirs. Indeed, the preponderance of studies – almost exclusively – have focused on estimating the influx *Q*(*t*) with very little attention paid to the expected residence time *T* other than to stipulate that it is greater than some perceived geologically relevant time horizon – usually 100 years. With respect to the main ocean carbon reservoirs, we can infer that up until recently (specifically, time scale of disturbance 75 years that is less than their residence times 500 years) they were in near equilibrium. That is *P* ≈*Q* from which Eq (1) follows.

The concept of carbon reservoirs up until now has only been applied to globally integrated processes. It can also be used to quantify much more targeted components of the biological pump and how they maintain sub-units of carbon reservoirs (Table 2). For instance (52) estimated the relative contribution to the overall DIC inventory following different export pathways (mixing pump, phyto-detritus, fecal pellets, vertical migrants) while (56) examined the contribution by various metazoan taxa (large, small, and mesopelagic fish, macro-, micro- and gelatinous zooplankton) and pathways (respiration, fecal pellets, deadfall) associated with the vertical migrants of Vinogradov ‘s ladder. These targeted reservoirs can be even more nuanced. For instance (57) estimated the respired DIC inventory for 5 species of overwintering copepods, (31) estimated the contribution from respired krill fecal pellets in the Southern Ocean, while (58) estimated the global contribution of cephalopod deadfalls. Dis-entangling the various pathways and populations engaged in maintaining the DIC reservoirs in the ocean provide a means of valuing

The concept of reservoir sub-units also exposes the flaw in an unqualified application of the CCS definition of sequestration to natural systems. We can, for instance, define a carbon reservoir as all respired carbon that has been in the ocean for more that *T*_*S*_ years. There is nothing in this definition that suggests that the associated carbon reservoir *B*_*S*_ is growing or contracting. Neither can a measurement the influx *Q*_*S*_ provide any information on this. Indeed, given the general condition of near equilibrium inferred from Holocene records, it is more likely that *B*_*S*_ = *Q*_*S*_*T*_*S*_ for any minimum time sequestration time scale. The only way to document sequestration ins a CCS sense, is to measure or predict changes in reservoir mass *dB*_*S*_*/dt*, or failing that, changes in turnover rates *dQ*_*S*_*/dt* and residence times *dT*_*S*_*/dt*.

### First passage and residence times

The main tool for making these estimates listed in Table 2 are transport matrix models (53, 62, 63) that distil the results of ocean circulation models, often tuned with biogeochemical data, to a matrix that describes the transport of tracer material from any box in a discretized ocean to any other box over a given time interval. The resulting matrix **A** is sparse in that many of its entries are zero – the probability of transport between distant locations in the ocean being vanishing small at typical model time scales of a year or so. The steady state reservoir size and mean residence time can be estimated as

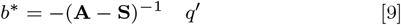

where *q*^*′*^(*x, y, z*) is the injection rate of DIC [gC m^−3^ yr^−1^] at a location (*x, y, z*) due to some process specific process (e.g. the respiration of a resident mesopelagic fish population), **S** is a strong sink term applied at the ocean surface, and *b*^∗^(*x, y, z*) is the spatial distribution of DIC [gC m^−3^] at steady state. Given *B*^∗^ = ∫_*V*_ *b*^∗^*DV* and *Q*^∗^=∫_*V*_ *q*′ *dV* as the integrate values over the entire volume of the oceans *V*, it follows that *T* ^∗^ = *B*^∗^*/Q* is the mean first passage time, a lower boundary to the mean residence time *T* . Further, since the first passage time (53) for the bulk of injected DIC is relatively short (i.e. the time between leaving the surface ocean as POC and being respired to DIC at depth; generally much less than 1 year), and accounting for the contribution of preformed DIC (3) discussed above, this suggests a rough heuristic that *B* = 1.6 *× B*^∗^ and correspondingly *T* = 1.6 *× T* ^∗^.

## ACKNOWLEDGMENTS

This work was supported by the Horizon Europe projects ECOTIP (869383), OCEAN-ICU (101083922), SEA-Quester (101136480), and by the Simons ‘ Foundation (931976).

## Notes

### Competing Interest Statement

The authors have declared no competing interest.

